# The clinician impact and financial cost to the NHS of litigation over pregabalin: a cohort study in English primary care

**DOI:** 10.1101/266403

**Authors:** Richard Croker, Darren Smyth, Alex J Walker, Ben Goldacre

**Author notes:** Dr Ben Goldacre (corresponding) Director, EBM DataLab Senior Clinical Research Fellow, Centre for Evidence Based Medicine Nuffield Department of Primary Care Health Sciences University of Oxford Radcliffe Observatory Quarter Woodstock Road Oxford OX2 6GG.

## Abstract

**Objectives:** Following litigation over pregabalin’s second-use medical patent for neuropathic pain NHS England were required by the court to instruct GPs to prescribe the branded form (Lyrica) for pain. Pfizer’s patent was found invalid in 2015; a ruling subject to ongoing appeals. If the Supreme Court appeal in February 2018 is unsuccessful, the NHS can reclaim excess prescribing costs. We set out to describe the variation in prescribing of pregabalin as branded Lyrica, geographically and over time; to determine how clinicians responded to the NHS England instruction to GPs; and to model excess costs to the NHS attributable to the legal judgments.

**Setting:** English primary care

**Participants:** English general practices

**Primary and secondary outcome measures:** Variation in prescribing of branded Lyrica across the country before and after the NHS England instruction, by practice and by Clinical Commissioning Group (CCG); excess prescribing costs.

**Results:** The proportion of pregabalin prescribed as Lyrica increased, from 0.3% over six months before the NHS England instruction (September 2014-February 2015) to 25.7% afterwards (April - September 2015). Although 70% of pregabalin is estimated to be for neuropathic pain, only 11.6% of practices prescribed Lyrica at this level; the median proportion prescribed as Lyrica was 8.8% (IQR 1.1-41.9%). If pregabalin had come entirely off patent in September 2015, and Pfizer had not appealed, we estimate the NHS would have spent £502m less on pregabalin to July 2017.

**Conclusion:** NHS England instructions to GPs regarding branded prescription of pregabalin were widely ignored, and have created much debate around clinical independence in prescribing. Protecting revenue from “skinny labels” will pose a challenge. If Pfizer’s final appeal on the patent is unsuccessful the NHS can seek reimbursement of excess pregabalin prescribing costs, potentially £502m.

### Strengths and weaknesses of the study

- We were able to measure the prescribing of pregabalin across all prescribing in England, eliminating bias.
- We were able to measure the impact of the NHS England’s instruction to GPs, which had a clear start and end date.
- Using the available data on prescribing volume, and changes to generic prices which occurred following the end of the patent, we were able to predict the excess costs to the NHS.
- The prescribing dataset does not include indication, therefore we were not able to ascertain whether individual patients given generic pregabalin after the NHS England letter were being treated for neuropathic pain; however branded prescribing rates were dramatically lower than estimates for the proportion of pregabalin prescribed for neuropathic pain.

## Abbreviations

CCG: Clinical Commissioning Group
GMC: General Medical Council
GP: General Practice
IQR: Interquartile range
NHS: National Health Service
NICE: National Institute for Health and Care Excellence
PCA: Prescription cost Analysis
QOF: Quality Outcomes Framework
SPC: Supplementary Protection Certificate

## Background

In August 2017 the price of pregabalin fell as it became fully generic. This was the end of a long process of legal claims, appeals, and unusual activity from NHS administrative staff, such as NHS England instructing all GPs and community pharmacies to use the branded form of the drug if it was being prescribed for neuropathic pain. With one last appeal outstanding, it is yet to be determined whether Pfizer were correct to claim exclusivity on this drug. As there are no known molecular or bioavailability differences with generic versions,[1] there is no clinical difference to using branded Lyrica.

This case raises a number of important issues to explore. The background of how drug patents operate; the value of generic prescribing; and NHS prescribing cost reimbursement are covered in Box 1. The basics of the Lyrica cases are covered in Box 2. We give more detail on the legal, ethical and financial issues raised by Lyrica in an accompanying Analysis paper. In this paper, we set out to describe whether NHS GPs followed the instructions given to them by NHS England; and the scale of the costs imposed on the NHS by the various ongoing legal cases.

#### Box 1. Generic prescribing, branded generics, and skinny labels.

##### Generic prescribing as a longstanding national policy

Prescribing all medication using the generic name of a drug is the near-universal norm throughout the UK, taught in all medical schools, and has had significant clinical and cost benefits: it provides a harmonised nomenclature which reduces complexity around prescribing, and so can reduce the risk of drug errors; generic names make it easy for clinicians and patients to know which class each drug belongs to; and it reduces costs as the least expensive drug is dispensed, with a switch to generic dispensing coming automatically at the end of a branded product’s patent life when cheaper generic forms enter the market.

##### Generics and “Branded Generics”

After a drug has lost its patent, other manufacturers may choose to market their version of the drug as a “branded generic”, i.e. a specific generic form of the drug with a distinct brand name. As an example in this case: Lyrica capsule 50mg is the “originator brand”; pregabalin capsule 50mg is the “generic”; Alzain capsule 50mg is a “branded generic”. When a generic is prescribed, the pharmacist can dispense any form of the drug: the originator brand, a branded generic, or any simple generic. When an originator brand is specified on the prescription, that specific originator brand must be supplied. When a branded generic is specified on the prescription, that specific branded generic must be supplied. Branded generics do not fall into the usual generic categories of the Drug Tariff, and are instead reimbursed at the specific price that the manufacturer has set; this is often lower that the originator brand or even the generic Drug Tariff price, to encourage use. Currently there are often cost saving opportunities from prescribing branded generics, and medicines optimisation teams commonly recommend these to their local GPs: however this comes at a cost of patients occasionally being changed to alternative presentations when prices change, resulting in possible confusion, concern, and reduced compliance (although this can also happen with generic prescribing, as the form dispensed can vary). Prescribing branded generics also breaches the widely supported general principle of doctors always prescribing the generic form.

##### Skinny Labels

A recent new challenge is the appearance of “skinny labels”: a marketing authorisation for branded or unbranded generics that permit them to be marketed for only a subset of all indications that the chemical is approved for, usually due to patent reasons. This is what has happened with pregabalin: pregabalin in the form of Lyrica has a marketing authorisation for use in anxiety, epilepsy, and pain; pregabalin as a generic only has a marketing authorisation for use in anxiety and epilepsy; if generic pregabalin is used for pain, then this is a “cross-label” use, outside of the marketing authorisation for the generic presentation, but (somewhat confusingly) within the marketing authorisation of the originator brand. This poses interesting new issues for debate, including for professional regulators: because the UK’s General Medical Council advice to prescribe within a treatment’s marketing authorisation is likely to have been drafted with patient safety in mind; but there are no known molecular or bioavailability differences between generic versions of pregabalin, only legal differences.

#### Box 2. Pregabalin Litigation

Pregabalin was originally protected by a product patent based on its antiseizure activity and expected efficacy for the treatment of, amongst other conditions, epilepsy and anxiety (the indications for which it was originally approved), which expired in May 2013. It was later found to be useful in the treatment of pain, and so a second “medical use” patent for this use was sought and granted, with an expiry date of 16th July 2017. The patent is actually in the name of Warner Lambert, a Pfizer subsidiary, but for convenience no distinction will be made here. In 2004 Pfizer was granted Supplementary Protection Certificates (SPCs) for pregabalin across Europe. SPCs are commonly granted to compensate for the period where a patent for a pharmaceutical product cannot be exploited because of the time taken to obtain regulatory approval for sale of the product. In this case, pregabalin for *all* indications would have been entirely exclusive to Pfizer until May 2018. However, the SPCs for pregabalin were never brought into force, as the necessary fees were never paid: it is currently not known why Pfizer chose not to activate the SPC. Consequently, when the data exclusivity period expired in 2014, then generic versions of pregabalin could be authorised and marketed for anxiety and epilepsy, but not for pain, as this continued to be covered by Pfizer’s “second medical use” patent.

Actavis applied for a marketing authorisation for their product (Lecaent), which had a “skinny label” covering only those indications no longer protected by patent. They were promptly sued for patent infringement by Pfizer, who demanded that Actavis ensured that Lecaent was not used for the treatment of pain. This is a problem, as manufacturers cannot easily monitor what generic products are supplied for: in the UK there is generally no indication written on the prescription, so the dispensing pharmacist does not know the reason for the prescription; in addition, UK doctors are trained and encouraged to prescribe drugs using the generic name, for a range of reasons including safety and cost effectiveness.

As an attempt to solve this problem, in 2015, NHS England were ordered by the Patents Court in London to issue instructions to Clinical Commissioning Groups, asking them to inform doctors that “Lyrica” should be specifically prescribed when pregabalin is used for neuropathic pain; and only prescribe pregabalin by its generic name when using the drug for other purposes. The NHS Business Services Authority was asked to instruct all community pharmacies that they should ensure, as far as possible, that Lyrica is dispensed where it was known that the prescription was for pain.

An additional complexity came in September 2015, when Pfizer’s own patent claim on neuropathic pain was itself held to be invalid, on the grounds that Pfizer had sought a patent for pain in general, but the data presented only plausibly supported efficacy against certain kinds of pain, including inflammatory pain. In relation to neuropathic pain, the data presented only supported peripheral neuropathic pain, not central neuropathic pain. Pfizer appealed, and again lost in the Court of Appeal in October 2016. Pfizer have appealed to the Supreme Court, and the case is programmed for February 2018. However, even if their claim to a patent on neuropathic pain is ultimately upheld, giving them some retrospective rights, the patent for neuropathic pain would have expired in July 2017.

## Methods

### Study design

We conducted a retrospective cohort study in English NHS GP practices: describing trends over time; measuring variation in the prescribing of branded Lyrica before and after the NHS England instruction; describing the impact of various legal actions on the total NHS expenditure on pregabalin; and modelling the additional costs to the NHS attributable to various legal judgements.

### Setting and data

We used data from our OpenPrescribing.net project. This imports openly accessible prescribing data from the monthly files published by the NHS Business Services Authority[2] which contain data on cost and volume prescribed for each drug, dose and preparation, for each month, for each English general practice. From this dataset we extracted data on all prescriptions dispensed between April 2013 and July 2016 for pregabalin capsules of any form.

### Impact of the court cases and NHS England Letter on pregabalin prescribing choices at a national level

We calculated the proportion of pregabalin capsules prescribed as generic, Lyrica, and other brands by using the 10th and 11th digits of the corresponding NHSBSA BNF code as either ‘AA’ (generic), ‘BB’ (Lyrica) or neither (other brands). This was plotted as a stacked bar chart. We present summary statistics on the change in prescribing practice.

### Impact of the NHS England Letter on practice-level prescribing choices

We included only practices that had a CCG code, and had at least 1,000 patients on list, to ensure we excluded prescribing in non-standard settings such as prisons or homeless services. From this list, we calculated the proportion of practices prescribing any Lyrica at all, and the average proportion of pregabalin that was Lyrica, per month, between September 2014-February 2015, and April - September 2015, which were the first six full months before and after NHS England issued the prescribing guidance. We present summary statistics on the change in prescribing practice.

### Variation between CCGs and practices in pregabalin prescribing choices

Using the CCG and practice data, we calculated CCG and practice level deciles for each month for the proportion of pregabalin prescribed as branded Lyrica, which were then plotted on a time series chart. We also used the OpenPrescribing.net measure “Pregabalin prescribed as Lyrica” to look for any variation in CCG implementation of the NHS England letter.[3]

### Modelling the impact of CCGs on Lyrica prescribing

In order to measure the extent to which practices’ prescribing behaviour was associated with their CCG we created a mixed effects linear regression model, with CCG as the only variable in the model, as a random effect. The outcome was Lyrica as a proportion of all pregabalin after NHS England issued their prescribing guidance to GPs (April - September 2015). The outcome was transformed using the Freeman-Tukey double arcsine transformation,[4] in order to transform the proportion while allowing zero values to be included. The model was used to calculate the significance of the association with CCG, as well as an R-squared value to describe the degree of variance associated with CCG membership.

### Modelling the cost to the NHS of the patent court cases

We calculated the cost to the NHS of the patent court case by applying the generic cost per capsule of August 2017 to the volume prescribed during the period Sept 2015 to Pfizer’s contested second medical use patent expiry date of 16th July 2017, and subtracting this from the actual NHS spend. The Drug Tariff price fell on the 1st August 2017. Our model assumes that, if the appeal had not continued after the September 2015 court case, then the reduction in generic cost price observed in August 2017 would have occurred on 1st October 2015; and that generic prescribing rates would have quickly moved back to the level before the NHS England guidance was released.

### Software and reproducibility

Data management and analyses were performed using Python, from a data warehouse in BigQuery, with additional data analyses performed using Stata 13.1. Data and all code are all available on Figshare.[5]

## Results

### Impact of the court cases on pregabalin prescribing choices at a national level

In the six months prior to the NHS England guidance being released, prescriptions for branded Lyrica were extremely rare, making up 6,944 out of 2,151,975 prescriptions (0.3%) for pregabalin (note that Lyrica would generally have been dispensed, as it was the only form of pregabalin on the market for most of this period). In the six months after the guidance was released, 604,450 prescriptions were for Lyrica, out of 2,350,450 pregabalin prescriptions in total (25.7%). From late 2015 there was also an increase in prescribing of other specific brands of pregabalin, which were less expensive than Lyrica. Generic pregabalin prescriptions continued to be reimbursed at the Lyrica cost, and therefore at the time these presented no financial benefit to the NHS, regardless of what brand the pharmacist dispensed (because for Category C drugs the reimbursement cost is set by what is prescribed rather than what is dispensed, and the generic pregabalin price during this period was based on Lyrica). During this period a number of CCGs formally added a branded generic to their prescribing formularies. Data on national prescribing is shown in Figure 1.

**Figure 1.**
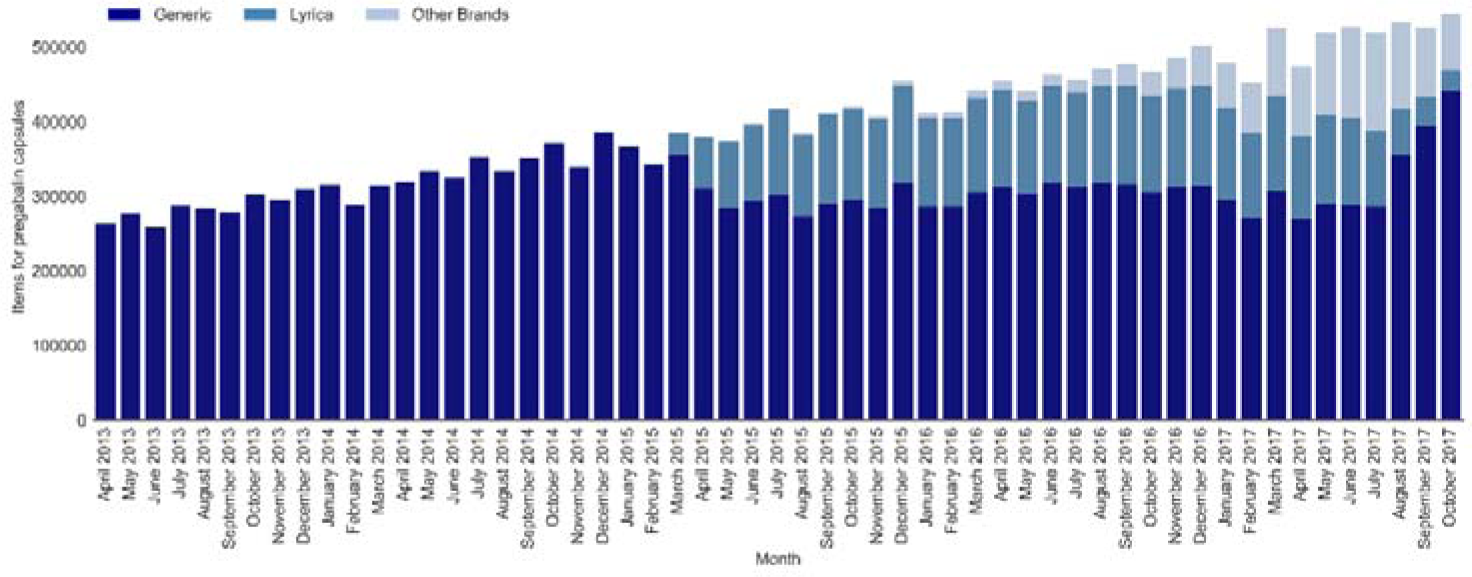
Volume of prescribing for generic pregabalin, Lyrica, and other brands, 2013-2017.

### Practice-level prescribing choices

While the national proportion of pregabalin prescribed as Lyrica increased from 0.3% to 25.7%, prescribing behaviour varied widely between practices, both before and after the NHS England letter. During the six months prior to the NHS England letter (September 2014 to February 2015) 898 out of 7,826 practices (11.5%) prescribed pregabalin as branded Lyrica at least once. However, even those that did prescribe the branded form did so very infrequently: among practices ever prescribing branded Lyrica, the branded form only accounted for 2.2% of all pregabalin prescriptions (mean 7.7 prescriptions for lyrica, and 341.9 for pregabalin).

In the first six months after the NHS England letter had been issued, the number of practices prescribing branded Lyrica at least once had increased to 6367 out of a total of 7736 prescribing pregabalin (82.3%). However, again, these practices tended to only use it for a minority of patients. Typically 70% of pregabalin prescriptions are thought to be for pain [6]. Only 11.6% of all practices met this 70% threshold after the NHS England letter; 78.6% of practices prescribed less than half their pregabalin as Lyrica; and 69.2% prescribed less than 30% as Lyrica. Overall after the NHS England letter 25.7% of pregabalin prescribed was Lyrica; but the median proportion of Lyrica prescribing at practice level was 8.8% (IQR 1.1%-41.9%). Therefore while most practices did change their prescribing to use Lyrica on occasion, most did so only marginally: NHS GPs in England largely ignored the direction given by NHS England to prescribe branded Lyrica.

The change in prescribing behaviour over time is presented in Figure 2, with deciles for the proportion of pregabalin prescribed as Lyrica by practice and by CCG. From this it can be seen that change came rapidly, but incompletely; that many practices prescribed no Lyrica; that most practices prescribed only a small amount of Lyrica compared to the 70% expected; and that prescribing behaviour began reverting to generic before the legal issues were resolved.

**Figure 2.**
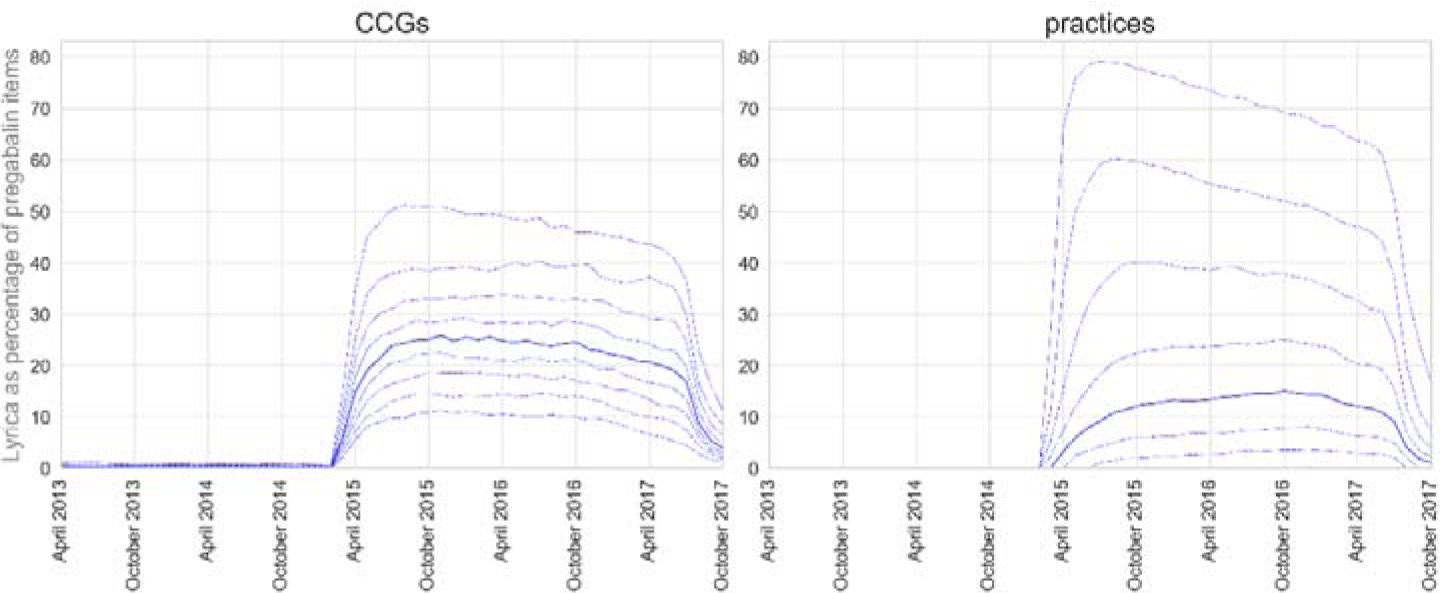
CCG and practice deciles for proportion of pregabalin capsules as Lyrica, 2013-2017.

### Modelling the impact of CCGs on Lyrica prescribing

At the CCG level, the proportion of Lyrica prescribing between April and September 2015 (after the guidance was issued) varied between 0.7% and 93.6% (median 21.2% IQR 13.5-31.7%). We assessed the extent to which a practice’s CCG membership was associated with the proportion of Lyrica prescribing, using the mixed effects model described above. This showed that there was a significant association with CCG membership (p<0.001), which was determined to account for 28.5% of the variance in the proportion of pregabalin prescribed as Lyrica.

The majority of CCGs followed the same pattern of change, albeit to varying levels, with a steep increase in the prescribing of Lyrica in April 2015, and then a corresponding decrease following the patent expiry in July 2017. However there were some exceptions where implementation did not follow this pattern. For example, Brent CCG had a relatively slow implementation of Lyrica prescribing, which only hit a peak shortly before the date of patent expiry (Figure 3). Ashford CCG appears to have implemented the letter quickly, but then the proportion of pregabalin prescribed as Lyrica started to reduce in early 2016, and was at below 10% by early 2017 (Figure 4). South Kent Coast CCG had an initial rapid implementation, and then a steady increase until late 2016, where the proportion of Lyrica reduced very quickly (Figure 5). The reasons for the variation in implementation are not clear, although in the case of reductions in Lyrica at Ashford and South Kent Coastal CCGs, similar profiles were also identified at Canterbury and Coastal, Swale, and Thanet, suggesting that there were collective discussions around how to manage regional implementation of the NHS England letter.

**Figure 3:**
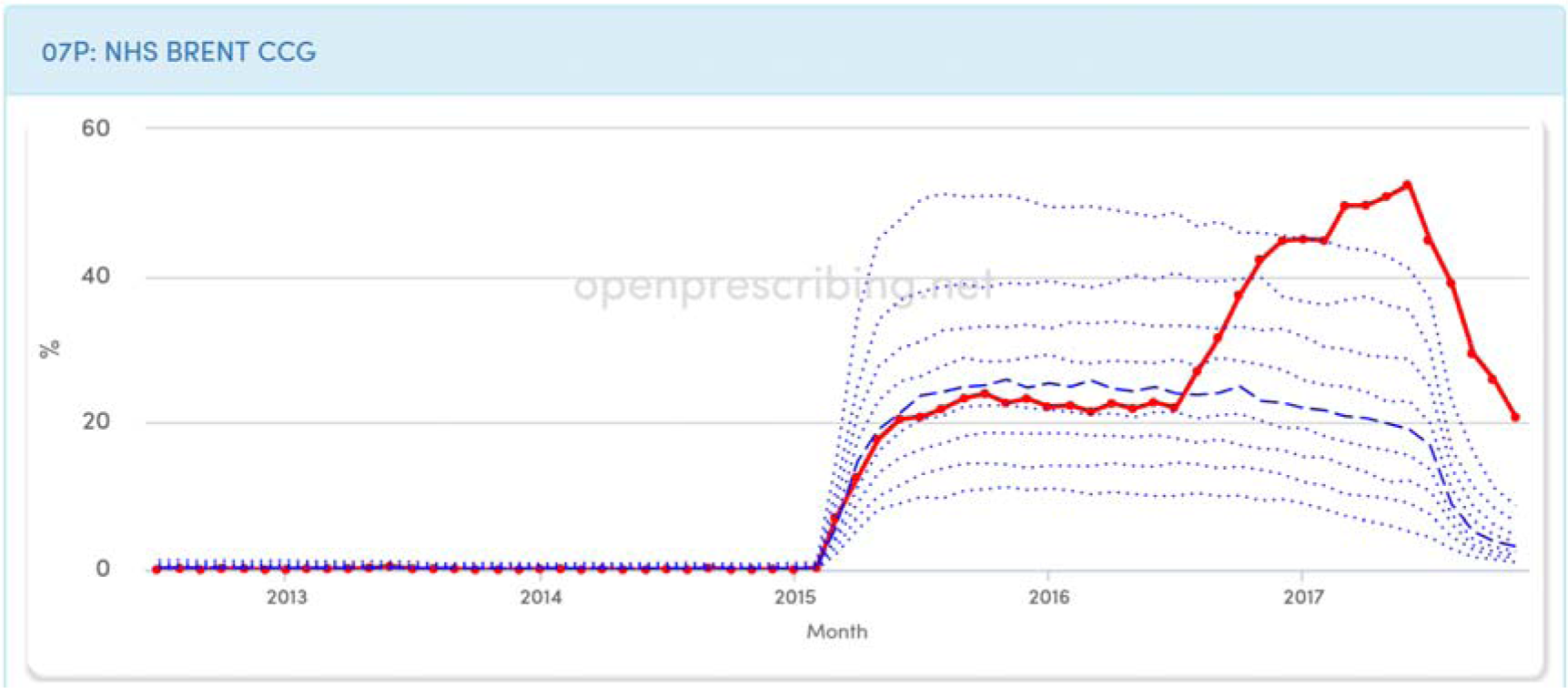
Brent CCG proportion of pregabalin capsules as Lyrica, 2012-2017.

**Figure 4:**
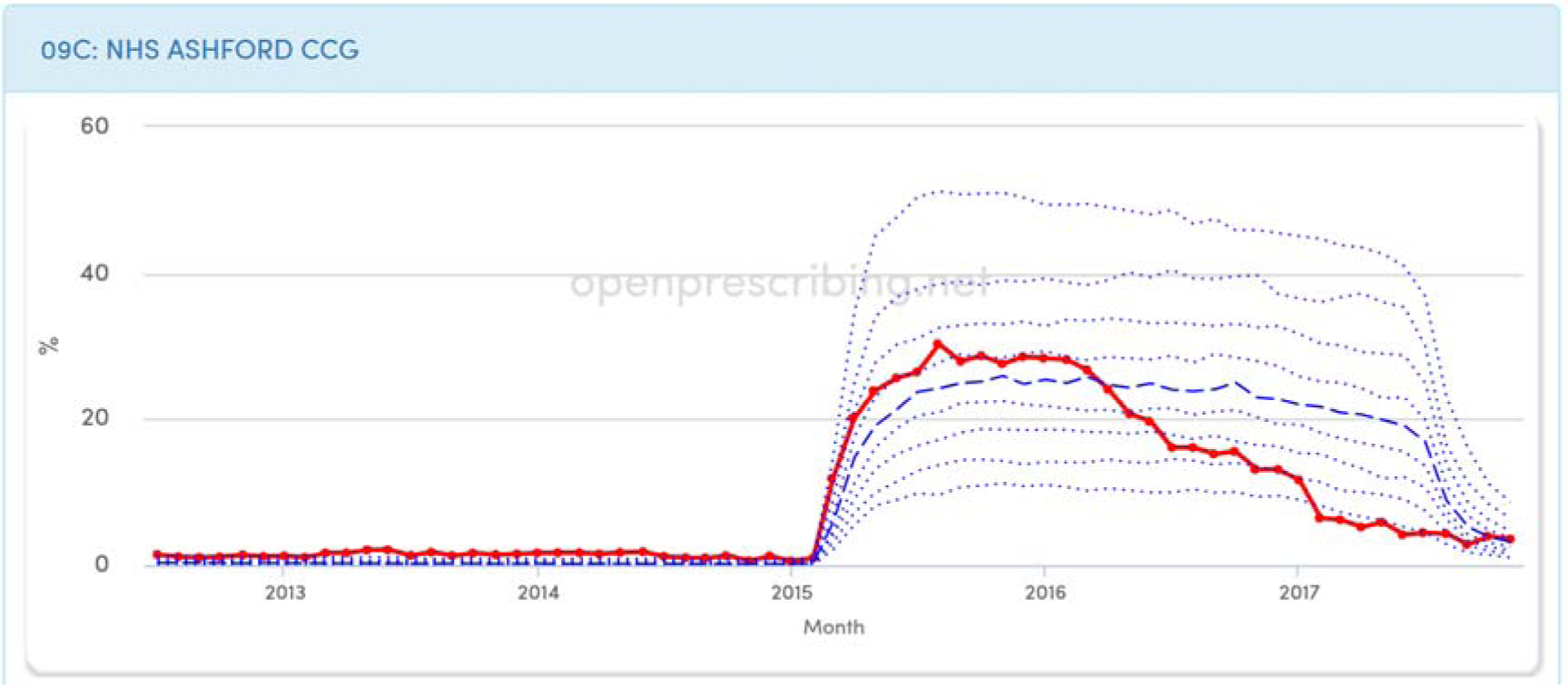
Ashford proportion of pregabalin capsules as Lyrica, 2012-2017.

**Figure 5:**
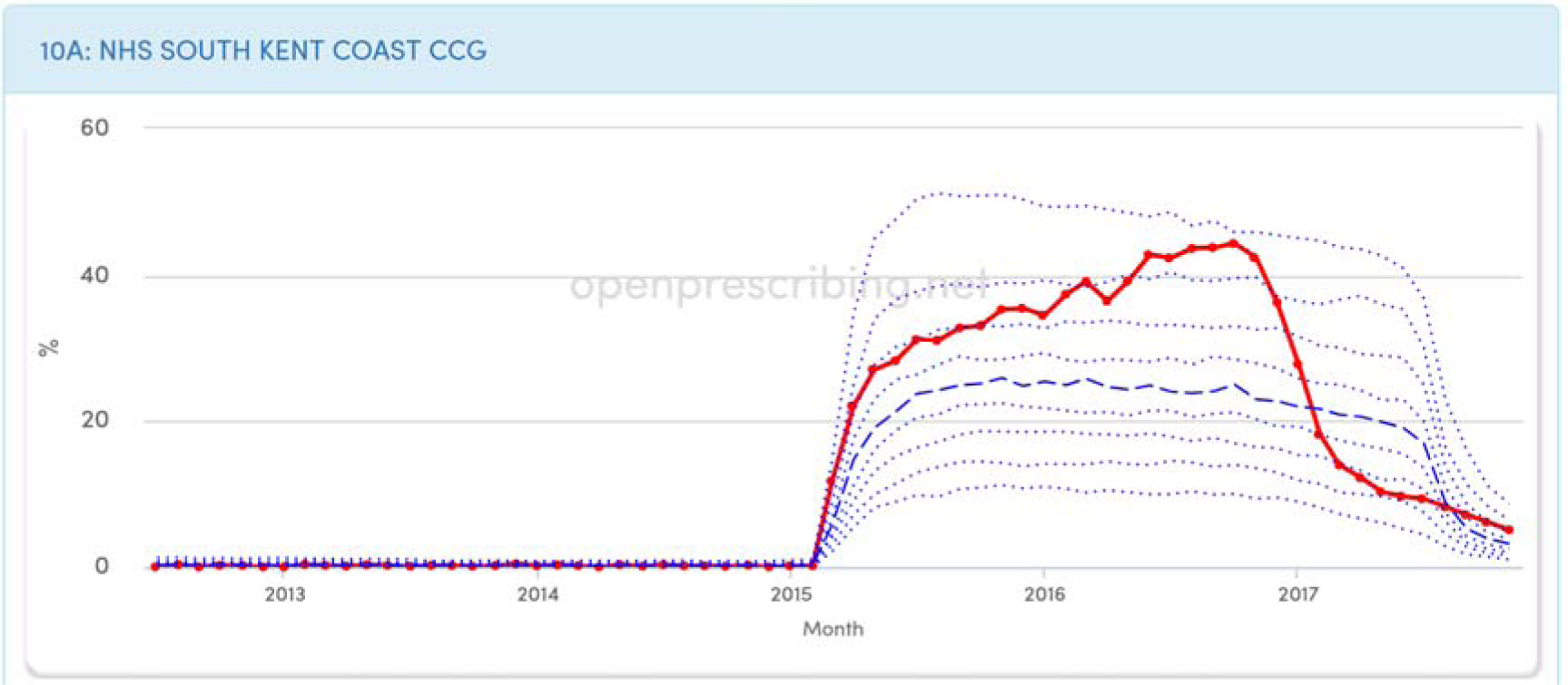
South Kent Coast CCG proportion of pregabalin capsules as Lyrica, 2012-2017.

### Modelling the cost to the NHS of the court case

The total net ingredient cost paid by the NHS for pregabalin capsules of varying strength during the period October 2015 to July 2017 was £564,095,548. The majority of this was prescribed either as generic pregabalin or branded Lyrica, for which the reimbursement price was the same. Our model, taking into account variation in price and prescribing volume for all strengths of pregabalin, calculates that the NHS would have paid £21,545,896 during this period if pregabalin had been available generically at the August 2017 drug tariff price, which is highly like to have been obtained had Pfizer not appealed against the adjudication on their patent not covering neuropathic pain. We therefore estimate an excess cost to the NHS of £501,912,683 attributable to the litigation over this patent that is currently in its final appeal. These calculations are presented, broken down by capsule strength, in Table 1. As with all prescribing expenditure it is possible that some of this excess cost attributable to the Lyrica patent would be affected by the Retained Margin process, used by the UK Department of Health to maintain community pharmacy profits at an agreed level (currently £800 million per year).[7]

**Table 1.** Estimates of excess cost to NHS, broken down by capsule.

| **Strength** | **NIC** | **Estimated**<br>**NIC at August**<br>**2017 Drug**<br>**Tariff** | **Difference in NIC** | **Difference in**<br>**Actual Cost** |
| --- | --- | --- | --- | --- |
| **100mg**<br>**capsules** | £65,935,836 | £2,104,684 | £63,831,151 | £59,050,198 |
| **150mg**<br>**capsules** | £110,670,782 | £4,070,097 | £106,600,685 | £98,616,294 |
| **200mg**<br>**capsules** | £34,042,634 | £1,318,460 | £32,724,174 | £30,273,133 |
| **225mg**<br>**capsules** | £12,959,623 | £653,483 | £12,306,141 | £11,384,411 |
| **25mg**<br>**capsules** | £54,732,394 | £1,651,112 | £53,081,282 | £49,105,494 |
| **300mg**<br>**capsules** | £89,364,123 | £6,008,264 | £83,355,859 | £77,112,505 |
| **50mg**<br>**capsules** | £78,998,270 | £1,930,269 | £77,068,001 | £71,295,608 |
| **75mg**<br>**capsules** | £120,226,538 | £3,917,799 | £116,308,739 | £107,597,214 |
| **Total** | £566,930,199 | £21,654,167 | £545,276,032 | £504,434,857 |
| **Total at**<br>**99.5%** | £564,095,548 | £21,545,896 | £542,549,652 | £501,912,683 |

## Discussion

### Summary

We found that NHS England guidance to prescribe branded Lyrica was widely ignored by NHS GPs in England: while most practices changed their prescribing behaviour to use Lyrica occasionally, most used it infrequently. We also found that CCG membership was a strong determinant of practices’ prescribing behaviour, accounting for 28.5% of the prescribing variance observed. The excess cost to the NHS attributable to Pfizer’s two appeals over Lyrica’s patent for neuropathic pain was estimated at approximately £502m. If the Supreme Court uphold the ruling, for a third time, that Pfizer’s second-use patent for neuropathic pain was invalid, then the NHS should take steps to recover this excess cost.

### Strengths and Weaknesses

We were able to use prescribing data to accurately show how doctors responded to NHS guidance. This data is complete, and covers the entire population of NHS English GPs, rather than a sample. It is likely to be a highly accurate record of what was dispensed, as it is the basis for NHS reimbursement to private pharmacies. As such, it represents the total dispensed, rather than prescribed; however there is no reason to believe patients are differentially less likely to present prescriptions for branded or generic pregabalin, therefore this is unlikely to be a source of bias. We are not able to account for variation between practices in the proportion of all pregabalin that was prescribed for pain, however previous work has shown that 70% of pregabalin prescribing is for pain, and the observed proportion was substantially short of this; while neuropathic pain is common, and unlikely to exhibit such non-uniform distribution as we have observed for Lyrica prescribing.

### Findings in Context

We are aware of one previous attempt in the grey literature to estimate costs of the Lyrica litigation, arguing that £54m could have been saved by the NHS between February and September 2015 through generic use; however this news piece gave no indication of how the figure was calculated, and in January 2016 the future generic price at launch in July 2017 could not have been known.[8]

### Policy Implications

The disputes around Lyrica raise several important issues. To our knowledge this is the first time a court has compelled the NHS to advise all doctors and pharmacists to change their usual prescribing and dispensing behaviour in order to ensure the protection of a drug patent. This may not be without unintended consequences: advising doctors to prescribe a branded medicine risks undermining the near universal advice that medicine should always be prescribed generically, itself a policy which has been highly successful in standardising prescribing behaviour and reducing costs.

Interestingly, our data shows that doctors largely did not do as NHS England instructed. Efforts to persuade doctors to use the Lyrica brand for neuropathic pain are likely to have been undermined by the fact that there was no medical reason to do so, as there is no evidence for a difference in clinical benefit between different manufacturers’ formulations of pregabalin; instead, there is a legal difference. Most doctors prescribed some Lyrica, but prescribed less frequently than would be expected if they were using it for neuropathic pain. It is possible that some doctors did so deliberately, prescribing some Lyrica to avoid being detected as “non-compliant”, while avoiding substantial changes to their prescribing choices. This would represent a form of civil disobedience. There has also been a degree of bad feeling towards Pfizer from the medical community as a consequence of their actions [9,10]. It will therefore be interesting to see whether the pharmaceutical industry undertakes similar action in the future.

Nonetheless, there is social value in patents being upheld, as this is the mechanism through which profits are used to incentivise innovation. Here, skinny labels represent a unique practical challenge. Pfizer were not able to capture the market share that their patent arguably warranted. It is hard to conceive of simple mechanisms to ensure differential reimbursement when the same drug is used for different purposes. Overhauling the entire prescription system to require indication is specified seems a heavy-handed response to a situation that has arisen only very rarely. Differences in formulation present another opportunity: doctors may have been more willing to use Pfizer’s form of pregabalin for neuropathic pain if it had been possible to create even a spurious distinction between two forms of the drug for two indications, such as different salts or doses for different indications.

Conversely, the NHS arguably overpaid for generic pregabalin that was prescribed for anxiety and epilepsy between 2015 and 2017, because the Category C Drug Tariff generic price was pegged to the list price of Lyrica, despite there being cheaper branded generics (such as Alzain and Axalid) available. Consequently pharmacists were reimbursed at this high Category C price for prescriptions written generically, regardless of what they dispensed, even if the prescription was clearly marked as being for epilepsy or anxiety, when a lower price would be justifiable. This is one of many areas where we believe the Drug Tariff requires reform in order to provide better value to the NHS.

The separate legal dispute over the legitimacy of the neuropathic pain patent itself poses a final challenge. Legal decisions - and a corporate body’s right to follow due process through the court, Court of Appeal and ultimately to the Supreme Court - can have large cost implications for the NHS. Our model shows that the NHS would have spent £502m less on pregabalin between October 2015 and July 2017, if the patent ruling had not been challenged by Pfizer. This equates to approximately 3.25% of the total NHS England primary care prescribing spend for that period. If Pfizer are unsuccessful in their final appeal, the NHS would be well served by seeking to retrieve these excess costs.

## Conclusions

The unprecedented court action regarding the prescribing of pregabalin was only partially successful, and has created much debate around the funding of drug research and intellectual property protection in a system where clinical independence to prescribe has been fundamental to practice in the NHS. If Pfizer is unable to defend its patent during the final appeal, the NHS could seek reimbursement of up to £502m.

## Acknowledgements

We are grateful to Helen Curtis for assistance with graphing techniques.

## Ethical approval

This study uses exclusively open, publicly available data; no ethical approval was required.

## Contributorship

RC and BG conceived the study. RC, BG and AW designed the methods. RC and AW collected and analysed the data with input from BG. RC, DS, AW and BG drafted the manuscript. All authors contributed to and approved the final manuscript. BG supervised the project and is guarantor.

## References

1 European Medicines Agency. Assessment Report: Pregabalin Mylan. 2015. http://www.ema.europa.eu/docs/en_GB/document_library/EPAR_-_Public_assessment_report/human/004078/WC500191300.pdf

2 Information Services Portal. NHS Business Services Authority. https://www.nhsbsa.nhs.uk/information-services-portal-isp (accessed 5 Feb 2018).

3 Pregabalin prescribed as Lyrica by all CCGs. OpenPrescribing. https://openprescribing.net/measure/lyrica/ (accessed 5 Feb 2018).

4 Freeman MF, Tukey JW. Transformations Related to the Angular and the Square Root. Ann Math Stat 1950; 21: 607–11.

5 Croker R, Walker A. The clinician impact and financial cost to the NHS of litigation over pregabalin. Figshare. https://figshare.com/projects/The_clinician_impact_and_financial_cost_to_the_NHS_of_litigation_over_pregabalin/29239 (accessed 14 Feb 2018).

6 Generics (UK) Ltd (t/a Mylan) v Warner-Lambert Company LLC [2015] EWHC 2548 (Pat). http://www.bailii.org/ew/cases/EWHC/Patents/2015/2548.html (accessed 28 Sep 2017).

7 Pharmaceutical Services Negotiating Committee. Margins survey. PSNC. https://psnc.org.uk/funding-and-statistics/pharmacy-funding/margins-survey/ (accessed 14 Feb 2018).

8 Price C. Pregabalin patent dispute could have cost the NHS up to £54m. Pulse Today. 2016. http://www.pulsetoday.co.uk/clinical/prescribing/pregabalin-patent-dispute-could-have-cost-the-nhs-up-to-54m/20030939.article (accessed 29 Sep 2017).

9 Jack A. Pfizer steps up battle to defend control of pregabalin. BMJ 2015; 350: h3119.

10 Jacobs M. GPs to shoulder costs of reversing patients from branded to generic pregabalin. Pulse Today. http://www.pulsetoday.co.uk/news/clinical-news/gps-to-shoulder-costs-of-reversing-patients-from-branded-to-generic-pregabalin/20035105.article (accessed 16 Oct 2017).

